# Depression and Anxiety in Healthcare Staffs in A Tertiary University Hospital in Luoyang, Henan Province: A Cross-Sectional Observational Study

**DOI:** 10.1101/516153

**Authors:** Hao Wang, Xiaochun Qing, Li Li, Lijun Zhang, Jing Ruan, Xiaohua Niu, Xiaotian Liu, Ruge Wu, Jiangbo Li, Keke Fan, Xuewei Chang, Mingming Zhang, Lixia Li, Zhuo Zhang, Shouyan Zhang

**Author notes:** Equal contribution (Hao Wang, Xiaochun Qing). **Corresponding author:** Dr. Shouyan Zhang, 288 Zhongzhouzhong Road, Luoyang, Henan Province, People’s Republic of China.

## Abstract

Mental health condition of medical professionals in China is under-recognized. The current study aimed to determine the prevalence of depression and anxiety among healthcare professionals and explore the potential influence factors.

**Method:** The study employed a cross-sectional design. All employees were surveyed in the first week of September 2017. General information included gender, age, workload, workplace violence, sleep quality and so on. Depression and anxiety were evaluated using PHQ-9 and GAD-7, respectively. SPSS 22.0 was used for data analysis. Logistic regression was conducted to explore risk factors contributed to metal health.

**Results:** A total of 1,950 questionnaires were delivered, and 1,864 were returned with a response rate of 95.6%. The prevalence of depression and anxiety were 24.1% and 28.9%. As for workload, the average number of beds in charge per month is 65.97±95.58 beds, among which internal medicine department and surgical department endure more workloads. Though workers in ER and ICU manage fewer beds, they bear the longest nonstop working length in the previous month (19.18±10.82 h). There were 78.0% had suffered WPV in the preceding year. Staffs in ER and ICU are at higher risk to physical violence, especially doctors. Regarding to sleep quality, only 9.2% participants reported that they sleep well. Logistic regression indicated that workplace violence and sleep quality were independent risk factors for both anxiety and depression.

**Conclusion:** The current study revealed the devastating conditions of mental disorders in medical workers and associated factors. Effective interventions are necessary to improve this situation.

## Introduction

Adverse mental health among medical workers not only impairs their own heath and life, but also hinders their job performance and reduces the quality of medical care patients received (1–3). Over a long period, it will bring great obstacles to the development of human resources in health fields. Due to the exposure to high levels of work-related stress, including extensive workload, long time of mental tension and frequent night shifts, physicians are more vulnerable to some mental disorders such as anxiety, depression, and occupational burnout than general population (4, 5). Anxiety and depression symptoms are the most common mental disorders. Previous studies conducted in United States, Britain, Norway and Japan, indicated that the prevalence of depressive symptoms among physicians ranged from 10% to 15% (6–9). Compared with medical professionals in foreign countries, Chinese healthcare givers encounter with some specific stressors. Firstly, Chinese doctors experience much greater pressure from work and living. Prior to the end of 2016, there were 3.19 million practicing (or assistant) physicians in China, amounting to 2.31 per thousand populations. Chinese Physicians dealt with 3.27 billion outpatient visits and 175.3 million inpatients in 2016 (10). 32.7% of doctors work more than 60 hours per week (11). What adds to this misfortune is that, their salary is much lower than peers overseas. According to the Physician Practice Report issued by the Chinese Medical Doctor Association in 2017, the average annual income of physicians was 75,000 yuan (about $11,220), in contrast to an average annual income of $299,000 for physicians in the United States (12, 13). What makes the situation worse is that the doctor-patient relationship in China is on the verge of collapse. Physicians and nurses not only are responsible for overloaded work, but also have to face the increasing threats to their personal safety at work (14). From 2009 to April 2015, there were more than 110 incidents resulting in severe injuries of clinical staffs in China (15). Although the issue of mental health condition of medical professionals in China has gradually been taken seriously, there are still few studies investigating the psychological distress of physicians in China. The current study aimed to identify the prevalence of depression and anxiety among Chinese physicians and evaluate potential risk factors that predispose them to psychological disorders.

## Method

### Study population

This study used a cross-sectional approach. An anonymous online questionnaire was conducted in a grade A tertiary hospital (Luoyang Central Hospital Affiliated to Zhengzhou University) in the city of Luoyang Henan Province (Central China) from 29 August 2017 to 7 September 2017. All subjects were employees in Luoyang Central Hospital Affiliated to Zhengzhou University. A total of 1950 questionnaires were distributed, of which 1864 valid questionnaires were retrieved (response rate = 95.6%).

### Questionnaire

Considering the limited and irregular break time available to clinical staffs, we decided to adopt a web-based questionnaire approach. A web page link to our questionnaire survey (https://www.wjx.cn/) was sent by mobile phone. The final questionnaire consisted of the following three sections.

#### 1. Socio-demographic information, lifestyles, work-related characteristics

Socio-demographic information included gender, age, education level, and marital status. Lifestyles items comprised of smoking status, drinking status and sleep quality. Work-related characteristics contained length of employment, work department, profession, number of beds in charge per month, the longest nonstop working time in the latest one month and workplace violence. Among these, subjective sleep quality was assessed using the Pittsburgh Sleep Quality Index (PSQI), which was developed by Dr. Buysse, a psychiatrist at the University of Pittsburgh in 1989 (16). There are 19 items in the scale. Each of them is assigned a score from 0 (no difficulty) to 3 (severe difficulty), with the total score ranging from 0 to 21 points calculated from seven components. This scale is not only validated for evaluating sleep quality in patients with sleep disorders, but also in healthy individuals. A score <5 is classified as good sleep quality and higher score reflects poorer sleep quality.

#### 2. Anxiety assessment

Anxiety assessment was completed using the seven-item generalized anxiety disorder scale (GAD-7). It is a brief self-questionnaire developed on the basis of the Diagnostic and Statistical Manual of Mental Disorders, fourth edition (DSM-IV) symptom criteria for GAD that allows for the rapid detection of GAD (17). The Chinese version of GAD-7 was downloaded on the Patient Health Questionnaire website (www.phqscreeners.com) (18). Subjects are required to answer whether, within the previous 2 weeks, they had experienced symptoms that tend to be associated with anxiety by answering seven items on a 4-point scale. The response options are: ‘‘not at all’’, ‘‘several days’’, ‘‘more than half the days’’, and ‘‘nearly every day’’; and they are scored 0, 1, 2, and 3, respectively. The total scores ranged from 0 to 21. Scores of 5, 10, and 15 represent cut points for mild, moderate, and severe anxiety, respectively. When used as a screening tool, the threshold score of 10 is recommended, at which the GAD-7 has a sensitivity of 89% and a specificity of 82% for GAD (19). So a score of 10 or higher is considered as anxiety in the following analysis.

#### 3. Depression assessment

We investigated depressive symptom using the nine-item Patient Health Questionnaire (PHQ-9), which is the depression module of the Primary Care Evaluation of Mental Disorders (PRIME-MD) and could also be used in general population, with high sensitivity and specificity (20). The PHQ-9 was translated into Chinese, and was downloadable on the Patient Health Questionnaire website (www.phqscreeners.com) (18). Participants are asked if they were bothered by depression related problems over the past two weeks. The possible answers and respective scores are ‘not at all’, ‘less that 1 week’, ‘1 week or more’, and ‘almost every day’. The answers to each question is scored on a scale of 0 to 3, hence with a total score of 0 to 27. Severity of depressive symptoms was reported as follows: none (score 1-4), mild (score 5-9), moderate (score 10-14), moderately severe (score 15-19), and severe (score 20-27). A cut point of 10 has been found to have high sensitivity (88%) and specificity (88%) for detecting depression (21, 22). So a score of 10 or higher is considered as depression in the following analysis.

### Ethical Approval

This study complied with Declaration of Helsinki and was approved by the Medical Ethics Committee of Luoyang Central Hospital before data collection commenced. Written informed consent could not be received due to the anonymous survey approach. Hence, oral informed consent for the survey was approved by the Medical Ethics Committee of Luoyang Central Hospital and obtained from every participant. Oral informed consent from participants in this research was obtained as follows: Firstly, our research team attended the morning shift meeting in every department and explained to all the employees. The explanations included the purposes, content and method of the survey, as well as the rights and privacy policies of the participants of the survey. And we specifically emphasized that, by submitting the answers, the participants were considered to automatically give the oral consent. Then, an anonymous questionnaire which was a two-dimension code easily to be transmitted via WeChat group was distributed via mobile phone.

### Statistical analysis

The data were analyzed using IBM SPSS version 22. Descriptive and frequency analyses were performed using mean, median, standard deviation and percentages. The chi-square test was used to compare the differences among two or more groups. Differences among groups were detected by parametric tests (t test, t’ test, ANOVA) when normality (and homogeneity of variance) assumptions are satisfied, otherwise the equivalent non-parametric test (Kruskal-Wallis H test) will be used. Odds ratios (ORs) and 95% confidence intervals (CIs) for each variable were calculated. Multivariable logistic regression analysis was used to screen the risk factors of anxiety and depressive (demographic variables, lifestyles and work-related characteristic variables as the independent variables). In this study, statistical significance was set at p<0.05 (two tailed).

## Results

### 1. Demographic characteristics

As shown in Table 1, a total of 1864 participants have been enrolled, with an average age of 32.49±8.17 years old. Among them, 20.4% are male, with an average age of 36.67±9.37 years old and 79.5% married; 79.6% are female, with an average age of 31.41±7.46 years old and 66.7% married. Most doctors are male (77.2%) and 98.4% have a bachelor’s degree (36.2%) or higher (62.2%) in terms of educational background. Majority of nurses are female (76.7%) and 52.5% have a bachelor’s degree (52.0%) or higher (0.5%). Only 5.1% of subjects are smokers and 8.3% consume alcohol, among which males and doctors occupy most.

**Table 1.**
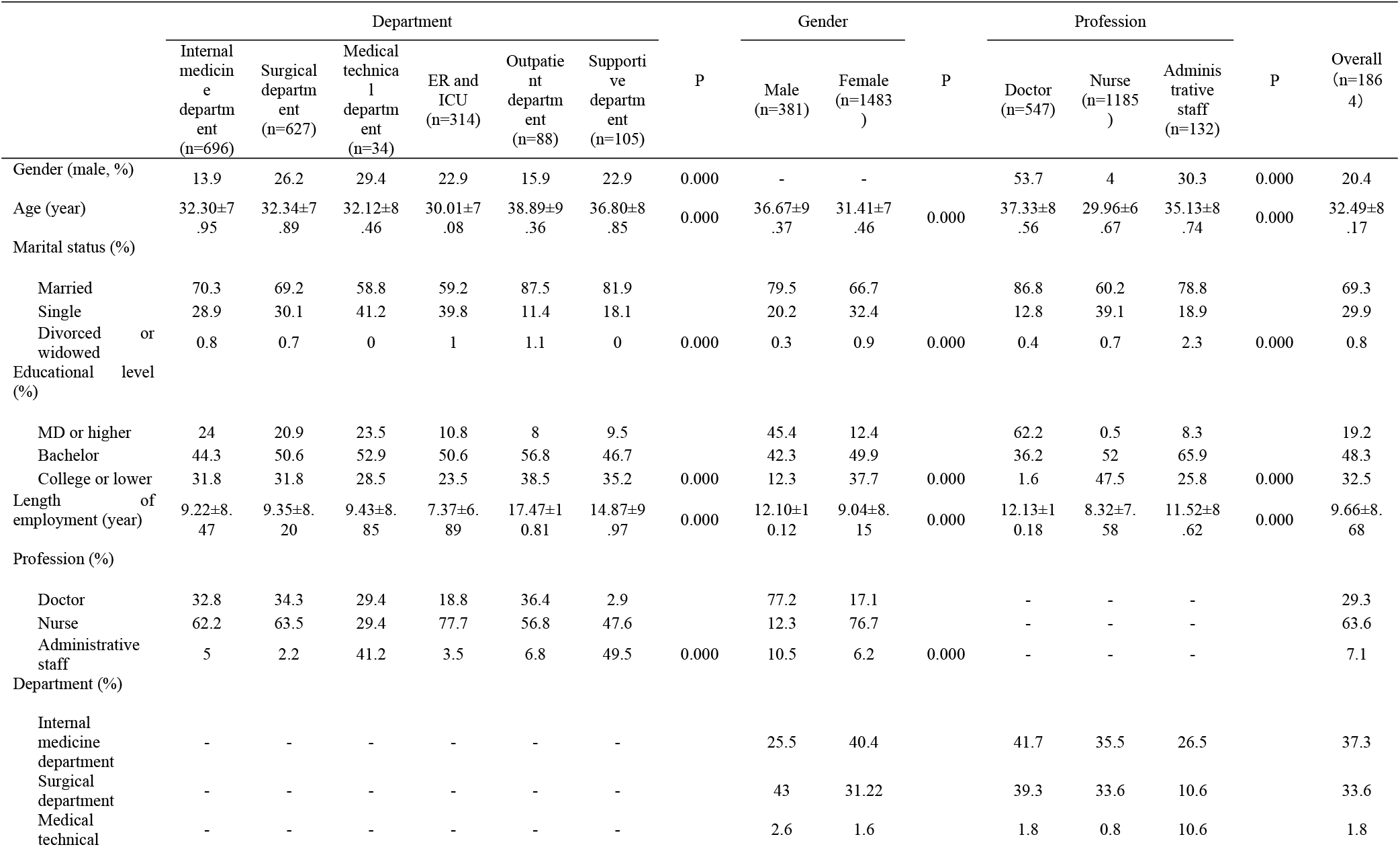
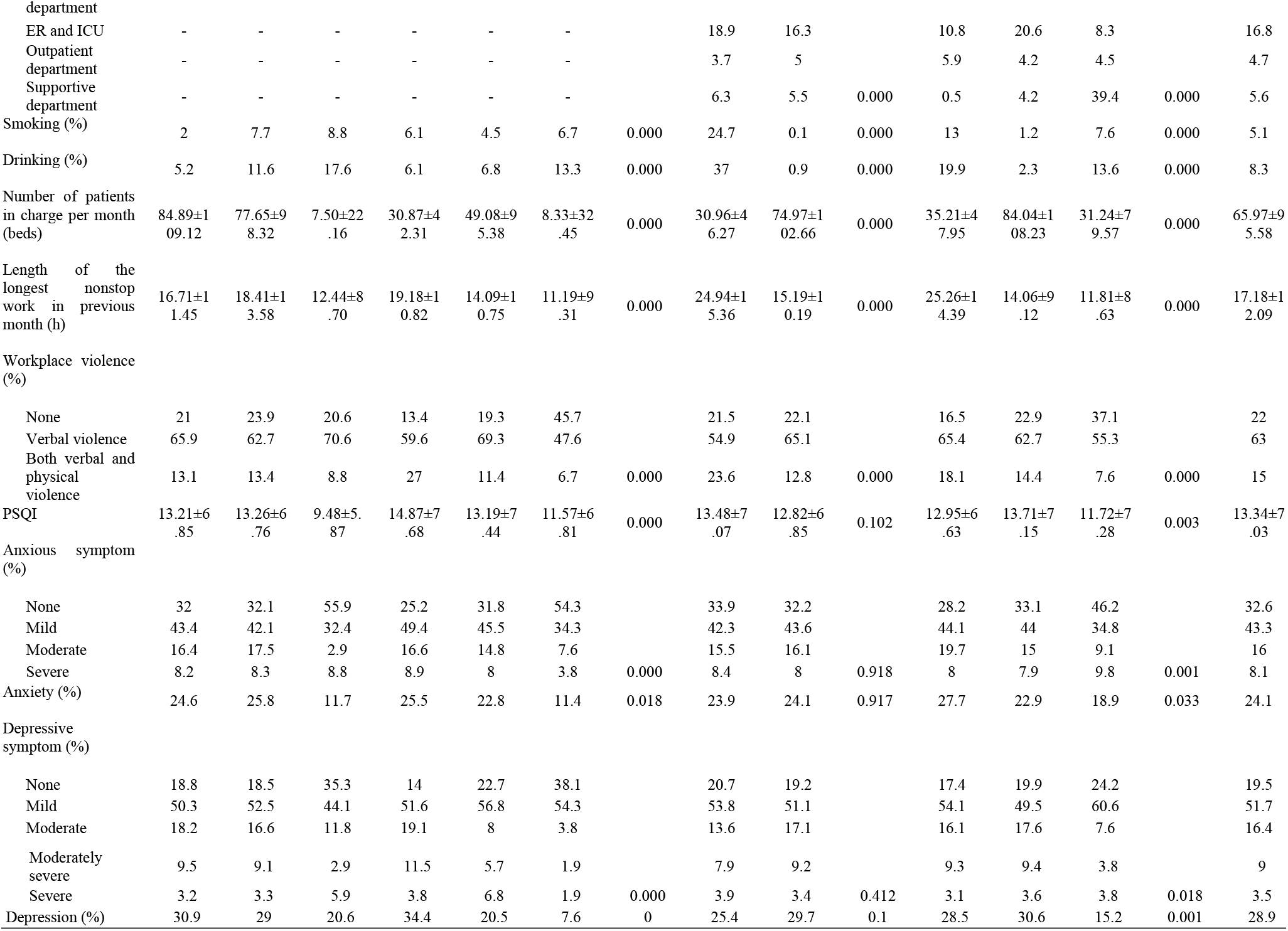
Demographic characteristics and prevalence of anxiety and depression.

### 2. Work-related information

The average length of employment is 9.66±8.68 years. Staffs in emergency room (ER) and intensive care unit (ICU) have the shortest time of employment (7.37±6.89 years), which might not only due to there are more nurses in these departments (77.7%), who tend to be relatively younger (29.96±6.67 years old) and with shorter experience length (8.32±7.58 years), but also because working in these departments means affording more work-related pressure, so that the employee turnover rate is higher. Outpatient department, by contrast, has the longest employment length (17.47±10.81 years), which may be owing to that outpatient service is mainly afforded by doctors with years of clinical experience. Personnel in supportive departments also have longer experience length, but majority of whom are not doctors, which is because human resources in these departments tend to be more stable due to relatively less working stress. The average number of beds in charge per month is 65.97±95.58 beds, among which internal medicine department and surgical department endure more workloads, who have to take charge of 84.89±109.12 beds and 77.65±98.32 beds, respectively, especially nurses. Though workers in ER and ICU manage fewer beds, they bear the longest nonstop working length in the previous month (19.18±10.82 h). Surgical department takes the second place with the nonstop working time of 18.41±13.58 h. Doctors (25.26±14.39 h) work longer than nurses (14.06±9.12 h) and administrative staffs (11.81±8.63 h).

With respect to exposure to violence at work, more than 78% of the participants have experienced workplace violence, including verbal violence or both verbal violence and physical violence. Personnel, especially physicians, in ER and ICU are at higher risk to physical violence, especially doctors. Regarding to sleep quality, only 9.2% participants reported that they sleep well. The sleep quality is the worst in ER and ICU. Doctors sleep better than nurses. No gender difference has been found in sleep quality.

### 3. Prevalence and severity of anxiety and depression

The evaluation results of anxiety and depression are also listed in Table 1. Overall, there are an estimated of 67.4% medical workers experience different extent of anxiety symptoms (mild: 43.3%, moderate: 16.0%, severe: 8.1%), 80.5% with depressive symptoms (mild: 51.7%, moderate: 16.4%, moderately severe: 9.0%, severe: 3.5%). Among them, 24.1% are screened as anxiety and 28.9% have depression. Internal medicine department, surgical department as well as ER and ICU have higher prevalence of anxiety, which occurrence rate of 24.6%, 25.8% and 25.5%, correspondingly. ER and ICU have the highest prevalence of depression (34.4%). In contrast, medical technical department and supportive department have the lowest prevalence of anxiety and depression. Doctors are slightly more susceptible to anxiety. No difference has been found between female and male.

### 4. Related risk factors for anxiety and depression

Associations between sample characteristics and anxiety or depression were assessed using logistic regression. As the characteristics included in the current study during the design phase were considered to be potential risk factors of anxiety and depression according to professional judgment and the findings of previous studies, the associations between each of them and anxiety or depression were screened using the logistic regression separately. The results were displayed in Figure 1 and Figure 2. Then, factors that reached statistical significances were included in the following multivariable logistic regression to explore for the independent risk factors. As shown in Figure 1, higher educational background (bachelor or higher), being doctor, clinical department, heavy workload, workplace violence (no matter verbal or physical) as well as poor sleep quality would increase the risk of anxiety, among which, educational level, nonstop work hours, workplace violence and sleep quality are independent risk factors for anxiety (Table 2). Similarly, the factors above, except educational background, would also increase the risk of depression (Figure 2). Department, violence and sleep quality are independent risk factors related to depression. However, age is an independent protective factor for depression (Table 3).

**Figure 1.**
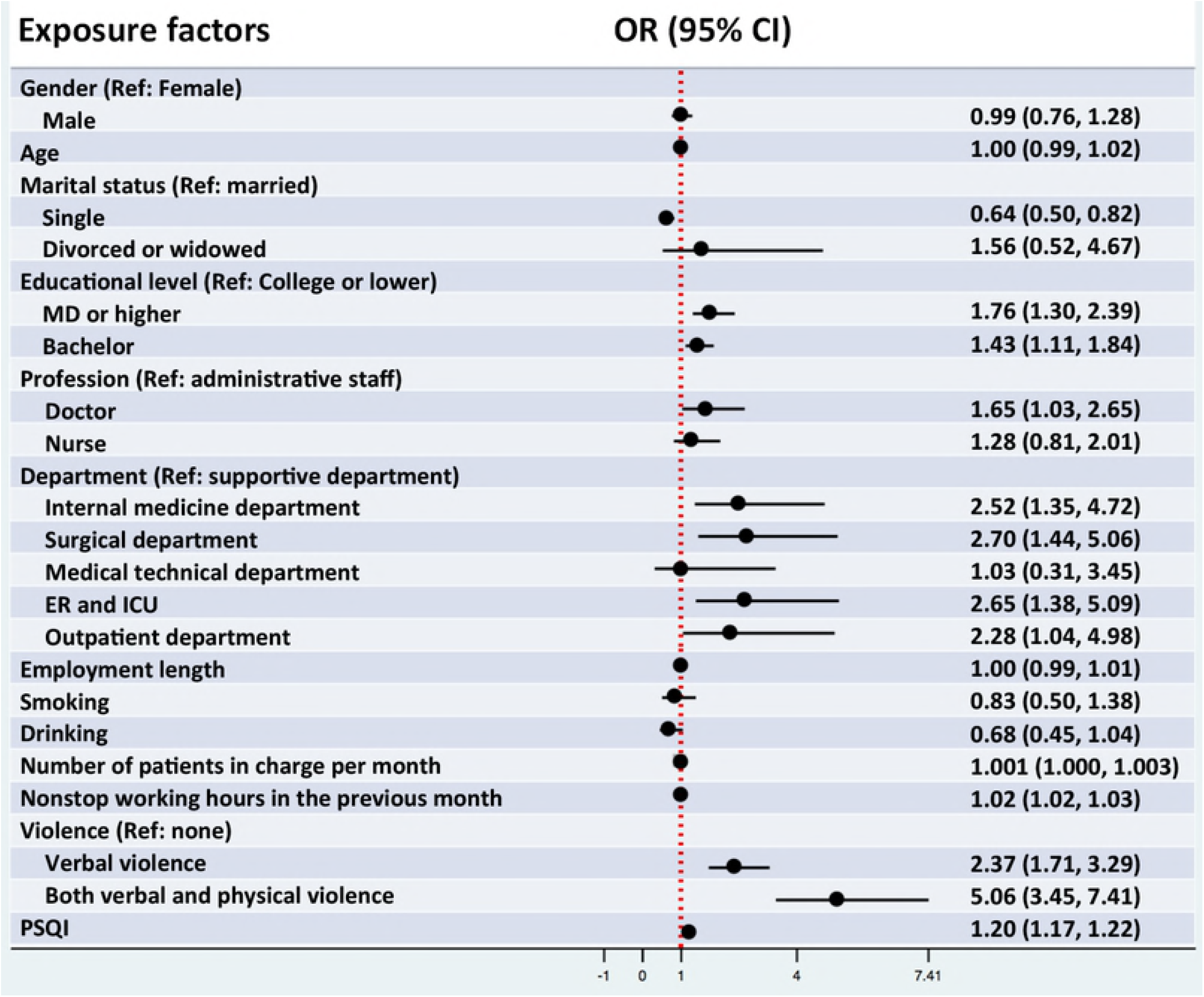
Association between sample characteristics and anxiety. The association between sample characteristics and anxiety was assessed using logistic regression analysis. It showed that higher educational background (bachelor or higher), being doctor, clinical department, heavy workload, workplace violence (no matter verbal or physical) as well as poor sleep quality would increase the risk of anxiety.

**Figure 2.**
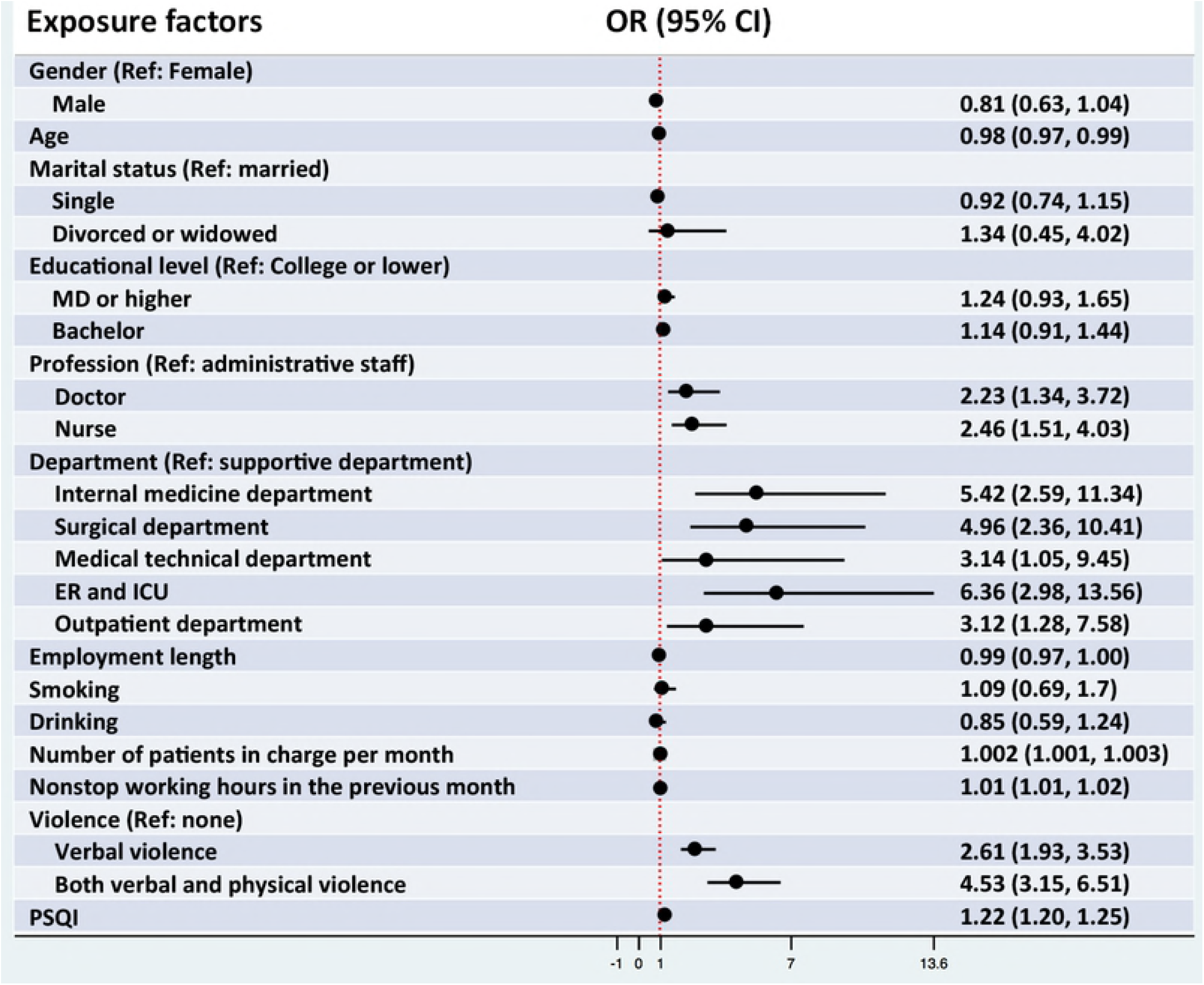
Association between sample characteristics and depression. The association between sample characteristics and depression was explored using logistic regression analysis. It indicated that being doctor, clinical department, heavy workload, workplace violence (no matter verbal or physical) as well as poor sleep quality would increase the risk of depression.

**Table 2.** Multivariable logistic regression for anxiety.

| Exposure factors | OR | 95% CI | p |
| --- | --- | --- | --- |
| Age (year) | 0.979 | [0.962, 0.995] | 0.012 |
| Department |  |  | 0.000 |
| Supportive department | 1.000 | reference |  |
| Internal medicine department | 6.163 | [2.657, 14.293] | 0.000 |
| Surgical department | 5.400 | [2.325, 12.542] | 0.000 |
| Medical technical department | 6.974 | [1.955, 24.872] | 0.003 |
| ER and ICU | 4.705 | [1.971, 11.232] | 0.000 |
| Outpatient department | 2.931 | [1.050, 8.186] | 0.040 |
| Violence |  |  | 0.001 |
| None | 1.000 | reference |  |
| Verbal violence | 1.602 | [1.129, 2.273] | 0.008 |
| Both verbal and physical violence | 2.326 | [1.501, 3.604] | 0.000 |
| PSQI | 1.224 | [1.197, 1.251] | 0.000 |

**Table 3.**
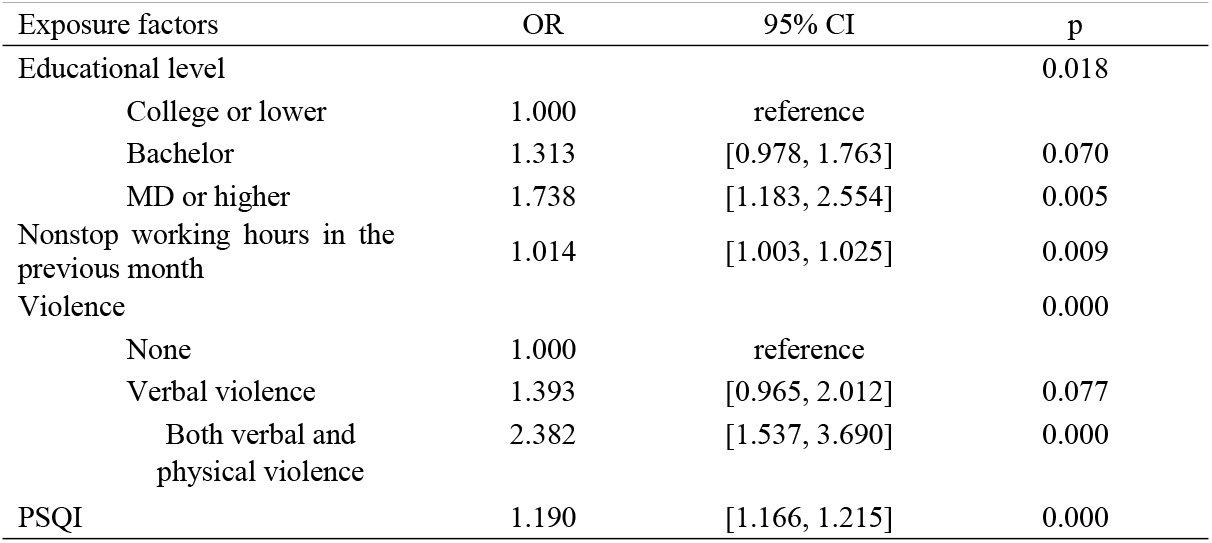
Multivariable logistic regression for depression.

## Discussion

In the current study, we found there are an estimated of 24.1% medical workers suffer from anxiety and 28.9% have depression, which is relatively higher than the results reported among physicians in other countries (6, 9). However, regarding to anxious and depressive symptom tendency, results obtained up to now differ from each other in China (23–25). In the current study, an estimated of 67.4% medical workers were found to have tendency of different extent of anxiety symptom (mild: 43.3%, moderate: 16.0%, severe: 8.1%), 80.5% with depressive symptom (mild: 51.7%, moderate: 16.4%, moderately severe: 9.0%, severe: 3.5%). While a study performed in Shenzhen public hospitals found that there were 25.67% and 28.13% physicians with anxious and depressive symptoms correspondingly (23). In Huiying Fang’s study performed among otorhinolaryngology nurses and physicians in Northern China, the data for depressive symptoms is as high as 57.2%(24). The data generated from public hospitals in Beijing is 37.1% for anxious or depressive symptoms (25). The differences among studies might be due to the following reasons. First, data reported in China generally exceed those obtained abroad, which might because of the different working environment between China and other countries, including heavier working load, lower income, and fragile doctor-patient relationship. Secondly, in China, work shift arrangement and workload also vary among hospitals with different grade or located in different cities. According to the National Health and Family Planning Commission of China, physicians working in tertiary hospitals have to deal with 8.1 outpatient visits and 2.7 beds per day; while, physicians at Community health care centers are responsible for dealing with more outpatient visits (15.9 visits per day) and less inpatients (0.6 beds per day) (10). Thirdly, the socio-demographic characteristics of participants enrolled in studies are also different, which would absolutely affect the final results. Lastly, the assessments for mental disorders used among studies are also different.

In the exploration of risk factors, we found that workplace violence is an independent risk factor for both anxiety and depression. Over the past few years, China has witnessed a surge in violence against medical professionals. Due to the possible violence which might come at any time and bring danger to their life, physicians have to afford excessive occupational stress, which absolutely will affect their psychological and physical health and then decrease their job performance (26–28). The current study also found that long duration of nonstop work and poor sleep quality were closely associated with psychological health, which was consistent with the conclusion arrived by previous studies (29–31). In the logistic regression for single values, it was indicated that working in clinical departments were more likely to be depressive, which might be due to the fast pace of work and the delicate doctor-patient relationship which may collapse at any time, especially in ER and ICU. Previous studies also found that doctors working in ER are much more susceptible to psychological disorders (32, 33). However, in multivariable regression, something interesting happened, a higher OR value was found in the medical technical departments than other departments. We did further examination of our data, and finally found that this abnormal OR value might result from the disparity in sample size among groups when limited to departments. So this result should be accepted with caution.

The current study revealed the severe station of mental disorders in medical workers and associated factors. This situation sometimes even affects the next generation of healthcare workers in China. In a survey among grade 3 to grade 7 medical students, 13.6% of the students clearly expressed their regrets of choosing the clinical medicine major (34). As for current physicians, about half of them said they intended to leave the profession, reported by a survey conducted in 29 public hospitals in Shandong province (35). And 64.48% of physicians responded that they would not encourage their child to become a physician in the future, according to the Chinese Medical Doctors’ Association (15). If left developing, Medicine may not be a favorite major among college applicants any more, and this situation will lead to great shortage of medical professionals (36). Given these negative effects, effective interventions are necessary to improve this situation.

### Limitations

There are some limitations of the current study. Firstly, we didn’t collect the information of income. Questions regarding income level did exist in the original questionnaire, but most workers refused to answer such private questions in our pilot study of the survey. Therefore, we removed this factor. Secondly, the factor of work shift is not included in our survey either. It is because the number and structure of employee group, as well as the work shift arrangement differ greatly among departments. What’s more, the definition of day shift and night shift are meaningless to surgeons, because most surgeons have to perform surgery even after an exhausted night shift ends. So we didn’t include factors such as work shift number per month. But, instead, we collected data of the number of beds in charge per month and the length of the longest nonstop working time in the latest month, which could also reflect the information of workload.

## Conclusion

The current study revealed that approximately one fifth of healthcare workers were suffering from anxiety and depression. Furthermore, workplace violence, poor sleep quality, and long work time were identified as the independent risk factors for psychological disorders. Policies, procedures and intervention strategies to improve this situation are required urgently.

## Acknowledgement

We are very grateful and thank all the colleagues participating in the current study from Luoyang Central Hospital Affiliated to Zhengzhou University. This work was financially supported by the Medical and Health Project of Luoyang Science and Technology Program (1820001A).

## Author Contributions

Conceptualization: SYZ HW.

Data curation: HW XCQ LL LJZ JR XHN KKF.

Formal analysis: HW XCQ XWC.

Funding acquisition: SYZ.

Investigation: HW XCQ LL LJZ JR XHN RGW JBL XTL KKF XWC LXL MMZ ZZ.

Methodology: HW XCQ JBL XTL.

Project administration: SYZ.

Resources: SYZ.

Supervision: SYZ.

Validation: SYZ LL.

Visualization: SYZ.

Writing ± original draft: HW.

Writing ± review & editing: SYZ HW XCQ.

## Reference

1. Tsai YC, Liu CH. Factors and symptoms associated with work stress and health-promoting lifestyles among hospital staff: a pilot study in Taiwan. BMC health services research. 2012;12:199.

2. Fahrenkopf AM, Sectish TC, Barger LK, Sharek PJ, Lewin D, Chiang VW, et al. Rates of medication errors among depressed and burnt out residents: prospective cohort study. BMJ (Clinical research ed). 2008;336(7642):488–91.

3. Ruitenburg MM, Frings-Dresen MH, Sluiter JK. The prevalence of common mental disorders among hospital physicians and their association with self-reported work ability: a cross-sectional study. BMC health services research. 2012;12:292–8.

4. Wallace JE. Mental health and stigma in the medical profession. Health (London, England : 1997). 2012;16(1):3–18.

5. Devi S. Doctors in distress. Lancet (London, England). 2011;377(9764):454–5.

6. Coomber S, Todd C, Park G, Baxter P, Firth-Cozens J, Shore S. Stress in UK intensive care unit doctors. British journal of anaesthesia. 2002;89(6):873–81.

7. Schwenk TL, Gorenflo DW, Leja LM. A survey on the impact of being depressed on the professional status and mental health care of physicians. The Journal of clinical psychiatry. 2008;69(4):617–20.

8. Vaglum P, Falkum E. Self-criticism, dependency and depressive symptoms in a nationwide sample of Norwegian physicians. Journal of affective disorders. 1999;52(1-3):153–9.

9. Wada K, Yoshikawa T, Goto T, Hirai A, Matsushima E, Nakashima Y, et al. Association of depression and suicidal ideation with unreasonable patient demands and complaints among Japanese physicians: a national cross-sectional survey. International journal of behavioral medicine. 2011;18(4):384–90.

10. National Health and Family Planning Commission of China. Statistical Communique on the Development of Health and Family Planning in China in 2016. 2016. [Available from: http://www.nhfpc.gov.cn/guihuaxxs/s10748/201708/d82fa7141696407abb4ef764f3edf095.shtml]

11. Wen J, Cheng Y, Hu X, Yuan P, Hao T, Shi Y. Workload, burnout, and medical mistakes among physicians in China: A cross-sectional study. Bioscience trends. 2016;10(1):27–33.

12. Chinese Medical Doctor Association. Chinese Physician Practice White Paper 2017. 2017. [Available from: http://www.cmda.net/u/cms/www/201807/06181247ffex.pdf.]

13. Medscape. Medscape Physician Compensation Report 2018. 2018. [Available from: https://www.medscape.com/slideshow/2018-compensation-overview-6009667?faf=1-2.]

14. Dalli J. Ending violence against doctors in China. Lancet (London, England). 2012;379(9828):1764.

15. Chinese Medical Doctor Association. Chinese Physician Practice White Paper 2015. 2015. [Available from: http://www.cmda.net/zlwqgzdt/596.jhtml.]

16. Buysse DJ, Reynolds CF, 3rd, Monk TH, Berman SR, Kupfer DJ. The Pittsburgh Sleep Quality Index: a new instrument for psychiatric practice and research. Psychiatry research. 1989;28(2):193–213.

17. Spitzer RL, Kroenke K, Williams JB, Lowe B. A brief measure for assessing generalized anxiety disorder: the GAD-7. Archives of internal medicine. 2006;166(10):1092–7.

18. Pfizer. Patient Health Questionnaire (PHQ) screeners [Available from: http://www.phqscreeners.com/]

19. Kroenke K, Spitzer RL, Williams JB, Monahan PO, Lowe B. Anxiety disorders in primary care: prevalence, impairment, comorbidity, and detection. Annals of internal medicine. 2007;146(5):317–25.

20. Santos IS, Tavares BF, Munhoz TN, Almeida LS, Silva NT, Tams BD, et al. [Sensitivity and specificity of the Patient Health Questionnaire-9 (PHQ-9) among adults from the general population]. Cadernos de saude publica. 2013;29(8):1533–43.

21. Kroenke K, Spitzer RL, Williams JB. The PHQ-9: validity of a brief depression severity measure. Journal of general internal medicine. 2001;16(9):606–13.

22. Manea L, Gilbody S, McMillan D. A diagnostic meta-analysis of the Patient Health Questionnaire-9 (PHQ-9) algorithm scoring method as a screen for depression. General hospital psychiatry. 2015;37(1):67–75.

23. Gong Y, Han T, Chen W, Dib HH, Yang G, Zhuang R, et al. Prevalence of anxiety and depressive symptoms and related risk factors among physicians in China: a cross-sectional study. PloS one. 2014;9(7):e103242.

24. Fang H, Zhao X, Yang H, Sun P, Li Y, Jiang K, et al. Depressive symptoms and workplace-violence-related risk factors among otorhinolaryngology nurses and physicians in Northern China: a cross-sectional study. BMJ open. 2018;8(1):e019514.

25. Xiao Y, Wang J, Chen S, Wu Z, Cai J, Weng Z, et al. Psychological distress, burnout level and job satisfaction in emergency medicine: A cross-sectional study of physicians in China. Emergency medicine Australasia : EMA. 2014;26(6):538–42.

26. Wu S, Lin S, Li H, Chai W, Zhang Q, Wu Y, et al. A study on workplace violence and its effect on quality of life among medical professionals in China. Archives of environmental & occupational health. 2014;69(2):81–8.

27. Jiang X, Gao J, Zhang X. Correlation analysis about workplace violence and job burnout among the head nurses. Chinese Nursing Management. 2008;8:18–20.

28. Zhao S, Xie F, Wang J, Shi Y, Zhang S, Han X, et al. Prevalence of Workplace Violence Against Chinese Nurses and Its Association with Mental Health: A Cross-sectional Survey. Archives of psychiatric nursing. 2018;32(2):242–7.

29. Zamanian Z, Nikeghbal K, Khajehnasiri F. Influence of Sleep on Quality of Life Among Hospital Nurses. Electronic physician. 2016;8(1):1811–6.

30. Wigg CM, Filgueiras A, Gomes Mda M. The relationship between sleep quality, depression, and anxiety in patients with epilepsy and suicidal ideation. Arquivos de neuropsiquiatria. 2014;72(5):344–8.

31. Lockley SW, Cronin JW, Evans EE, Cade BE, Lee CJ, Landrigan CP, et al. Effect of reducing interns’ weekly work hours on sleep and attentional failures. The New England journal of medicine. 2004;351(18):1829–37.

32. Yahaya SN, Wahab SFA, Yusoff MSB, Yasin MAM, Rahman MAA. Prevalence and associated factors of stress, anxiety and depression among emergency medical officers in Malaysian hospitals. World journal of emergency medicine. 2018;9(3):178–86.

33. Alharthy N, Alrajeh OA, Almutairi M, Alhajri A. Assessment of Anxiety Level of Emergency Health-care Workers by Generalized Anxiety Disorder-7 Tool. International journal of applied & basic medical research. 2017;7(3):150–4.

34. Li D, Yin W, Zhang X, Su M, Meng M, Wang Q. Investigation on turnover intention of medical staff in public hospitals and research of early warning system’s construction. Chin J Hosp Adm. 2010;26:218–21.

35. Han X, Wang Y, Zhao J, Pan H, Yu J. Examining influence of violence against physicians on Chinese medical students’ career choice. Chinese medical journal. 2014;127(24):4287–9.

36. Song P, Jin C, Tang W. New medical education reform in China: Towards healthy China 2030. Bioscience trends. 2017;11(4):36.

